# Sensory Eye Dominance Plasticity in the Human Adult Visual Cortex

**DOI:** 10.1101/2023.04.21.537873

**Authors:** Ka Yee Kam, Dorita H. F. Chang

**Affiliations:** Department of Psychology, The University of Hong Kong, Hong Kong; The State Key Laboratory of Brain and Cognitive Sciences, The University of Hong Kong, Hong Kong

**Keywords:** Sensory eye dominance, dichoptic perceptual training, perceptual learning, plasticity, fMRI

## Abstract

Sensory eye dominance occurs when the visual cortex weighs one eye’s data more heavily than those of the other. Encouragingly, mechanisms underlying sensory eye dominance in human adults retain a certain degree of plasticity. Notably, perceptual training using dichoptically presented motion signal-noise stimuli has been shown to elicit changes in sensory eye dominance both in visually impaired and normal observers. However, the neural mechanisms underlying these learning-driven improvements are not well understood. Here, we measured changes in fMRI responses before and after a five-day visual training protocol to determine the neuroplastic changes along the visual cascade. Fifty visually normal observers received training on a dichoptic or binocular variant of a signal-in-noise (left-right) motion discrimination task over five consecutive days. We show significant shifts in sensory eye dominance following training, but only for those who received dichoptic training. Pattern analysis of fMRI responses revealed that responses of V1 and hMT+ predicted sensory eye dominance for both groups, but only before training. After dichoptic (but not binocular) visual training, responses of V1 and hMT+ could no longer predict sensory eye dominance. Our data suggest that perceptual training-driven changes in eye dominance are driven by a reweighting of the two eyes’ data in both primary and task-related extrastriate visual areas. These findings may provide insight into developing region-targeted rehabilitative paradigms for the visually impaired, particularly those with severe binocular imbalance.

## Introduction

Information from the two eyes is not necessarily weighed equally at the level where it is integrated. Such functional asymmetry of the two eyes, thought to originate from the visual cortex is known as *sensory eye dominance*. Sensory eye dominance is a prominent characteristic exhibited by some clinical populations, such as patients with strabismus amblyopia (Ding et al., 2013; Zhou et al., 2013), but is also observed in the normal population. Previous studies have found that around 60% of the healthy population shows mild dominance while a significant minority (30-40%) shows strong dominance (Li et al., 2010; Yang et al., 2010; Zhang et al., 2011). Of immediate relevance to the present study, a growing body of work has indicated that mechanisms underlying sensory eye dominance in human adults retain a certain degree of plasticity. In particular, a visual training protocol using dichoptically presented signal-in-noise motion stimuli has gained special traction as it has been demonstrated to effectively reduce eye dominance in both the visually impaired (Hess et al., 2010) and normal observers (Kam and Chang, 2021). To date, much of the research has been focused on developing different paradigms to promote eye-rebalancing (Li et al., 2013; To et al., 2011; Tuna et al., 2020; Xu et al., 2012, 2010), but very little work has been done to reveal the neural underpinnings of sensory eye dominance and its plasticity. Knowledge of such mechanisms may provide insight into how one might develop an efficient rehabilitative protocol.

While little remains known as to the exact neural substrates underlying sensory eye dominance, speculative models of how eye imbalance may arrive have been put forth. The two-stage model for binocular integration highlights that the visual system adjusts the relative strength of each eye’s input and integrates these inputs in two distinct stages: The first stage of contrast gain control occurs before binocular combination, where the data from the two eyes remain segregated, but each eye receives inhibitory input from the contralateral eye while the second stage of gain control occurs after binocular combination (Meese et al., 2006). Theoretically, learning-related changes in sensory eye dominance could arise from the changes before, at, or after binocular combination. While this model provides a clear theoretical basis for the modulation and integration of the two eyes’ inputs under normal conditions, the sites of the two posited stages (pre- and post-binocular summation) are still largely unknown. To probe the potential loci of eye balance plasticity, we previously introduced four dichoptic training tasks that differed in terms of the presence of external noise and the visual feature implicated and examined their capacity to drive changes in sensory eye dominance (Kam and Chang, 2021). We found that changes in sensory eye dominance do not depend on the trained task or visual feature, suggesting that the dichoptic training paradigm may at least partially act to balance interocular suppression before or at the site of binocular combination. Therefore, there is reason to believe that dichoptic visual training may act on mechanisms early in the visual cascade, potentially in the lateral geniculate nucleus (LGN) or the primary visual cortex (V1).

The lateral geniculate nucleus is the primary source of feedforward input to V1 (Hendrickson et al., 1978; Hubel and Wiesel, 1972), but it also receives a vast amount of descending feedback from the visual cortex (Van Horn et al., 2000). While LGN neurons are monocular, animal studies have demonstrated that there are interocular inhibitory interactions in the LGN (Dougherty et al., 2021; Marrocco and McClurkin, 1979; Rodieck and Dreher, 1979; Sanderson et al., 1971; Sengpiel et al., 1995; Xue et al., 1987) mediated either by local interthalamic circuits (Guillery and Colonnier, 1970) or corticogeniculate feedback from V1 (Freeman and Tsumoto, 1983; Gaska et al., 2000). The cortico-geniculate feedback projection originating from V1 and (indirectly from) hMT+ is speculated to modulate the strength of the two eyes’ signals before binocular combination (Dougherty et al., 2019b, 2021). It is, therefore, possible for the two eyes to exert different strengths of gain control over signals coming from the other eye at the point where the two eyes’ data are still segregated. While different LGN responses to inputs from one eye or another are not well reported, one fMRI study that used high contrast checkerboard stimuli to assess the functional integrity of the LGN in human amblyopia reported lower LGN activation when with inputs from the amblyopic versus the fellow eye (Hess et al., 2009). Moreover, a recent diffusion-weighted imaging study revealed that the white matter microstructural properties of the optic radiations are able to predict the magnitude of sensory eye dominance in the visually normal adults (Chan and Chang, 2022). These findings altogether suggest that mechanisms early in the visual cascade, perhaps the speculative interocular gain control mechanisms at the LGN, may underlie sensory eye dominance in normal-sighted individuals.

To our knowledge, changes of eye-specific responses in the LGN has only been demonstrated in rodents that have been monocularly deprived for a week (Jaepel et al., 2017). A recent functional brain imaging (fMRI) study involving visually healthy human adults did not observe changes in LGN after 2-hours of monocular deprivation (Kurzawski et al., 2022). It is worth noting, however that the neural mechanisms underlying this kind of short-term plasticity may be different from those induced by a more extended period of monocular deprivation (Ramamurthy and Blaser, 2021) or binocular perceptual training. Here, then, we deem LGN as a particular site of interest to probe changes following dichoptic perceptual training.

We also considered primary visual cortex (V1) as another potential site of interest. From the classic work of Hubel and Weisel (Hubel and Wiesel, 1968a), layer three of V1 is a potential locus of binocular combination as it is the layer within which binocularity emerges. Drawing upon data from more recent work on normal vision, V1 showed an asymmetrical trend of activation during monocular stimulation: higher fMRI signal magnitude during stimulation of the dominant eye and lower magnitude during stimulation of the non-dominant eye (Conner et al., 2007). Studies using visual evoked potential (VEP) (Lunghi et al., 2015a) and fMRI (Binda et al., 2018) have shown that following short-term (2-hour) monocular deprivation, activity in V1 increases for the deprived eye but decreases for the non-deprived eye. Further, the activity changes in V1 are correlated with changes in eye dominance. It is unclear, however, as to whether activity changes in V1 might similarly come about after longer-term dichoptic visual training – or whether long-term training results in a different set of neural changes altogether.

The third site we deemed interesting to probe with dichoptic visual training is the human middle temporal complex (hMT+). It is well-established from classical neurophysiological studies (Beckers and Hömberg, 1992; Braddick et al., 2001; Vaina et al., 2005) that hMT+ is involved in motion processing. As the popular dichoptic training task involves a motion-stimulus, hMT+ then could be a possible stimulus-specific locus for the post-summative improvements, although it has not been shown to link specifically to sensory eye dominance plasticity.

Here, using fMRI, we aimed to identify the neural changes associated with improvements in sensory eye balance as driven by dichoptic visual training. We measured changes in blood oxygenated level-dependent (BOLD) activity before and after training in our main sites of interest. We contrasted learning effects following dichoptic signal-in-noise motion training (signal and noise dots presented to different eyes) with training on a binocular variant of the same task (signal and noise dots presented to both eyes). We elected to include a binocular variant of the task for training as it ensures that any differences in learning (and learning associated neural changes) would be due solely to the mode of presentation (dichoptic vs binocular). Based on previous work, we predicted that training on the dichoptic, but not binocular variant of the motion signal-noise task would result in behavioural shifts in eye dominance. Further, we reasoned that if perceptual training using dichoptically presented signal-in-noise motion stimuli affects pre-binocular-summation mechanisms, we would observe training-related changes early in the visual cascade (i.e., in the LGN), perhaps through cortico-geniculate feedback. By contrast, if dichoptic perceptual training acts on mechanisms at or immediately after binocular summation, we would observe learning-associated changes in V1. Lastly, it may well be that dichoptic training results in a reweighing of the signals in higher-order visual mechanisms that are training-feature-specific (in this case, motion-established hMT+).

## Results

### Behaviour

First, we examined the degree of learning achieved by comparing the performance during “early” versus “late” training blocks. For each subject, we averaged thresholds obtained from the first three training blocks and the last three training blocks to represent task performance of the early and late training stages, respectively. We then performed (corrected) paired t-tests independently for the two training groups (Figure 2a & b). The analyses indicated significant improvements in the trained task (i.e. lower thresholds for late vs early training) for both the dichoptically (t_(23)_ = 5.616, p < .001) and binocularly (t_(24)_ = 4.158, p < .001) trained groups.

**Figure 1.**
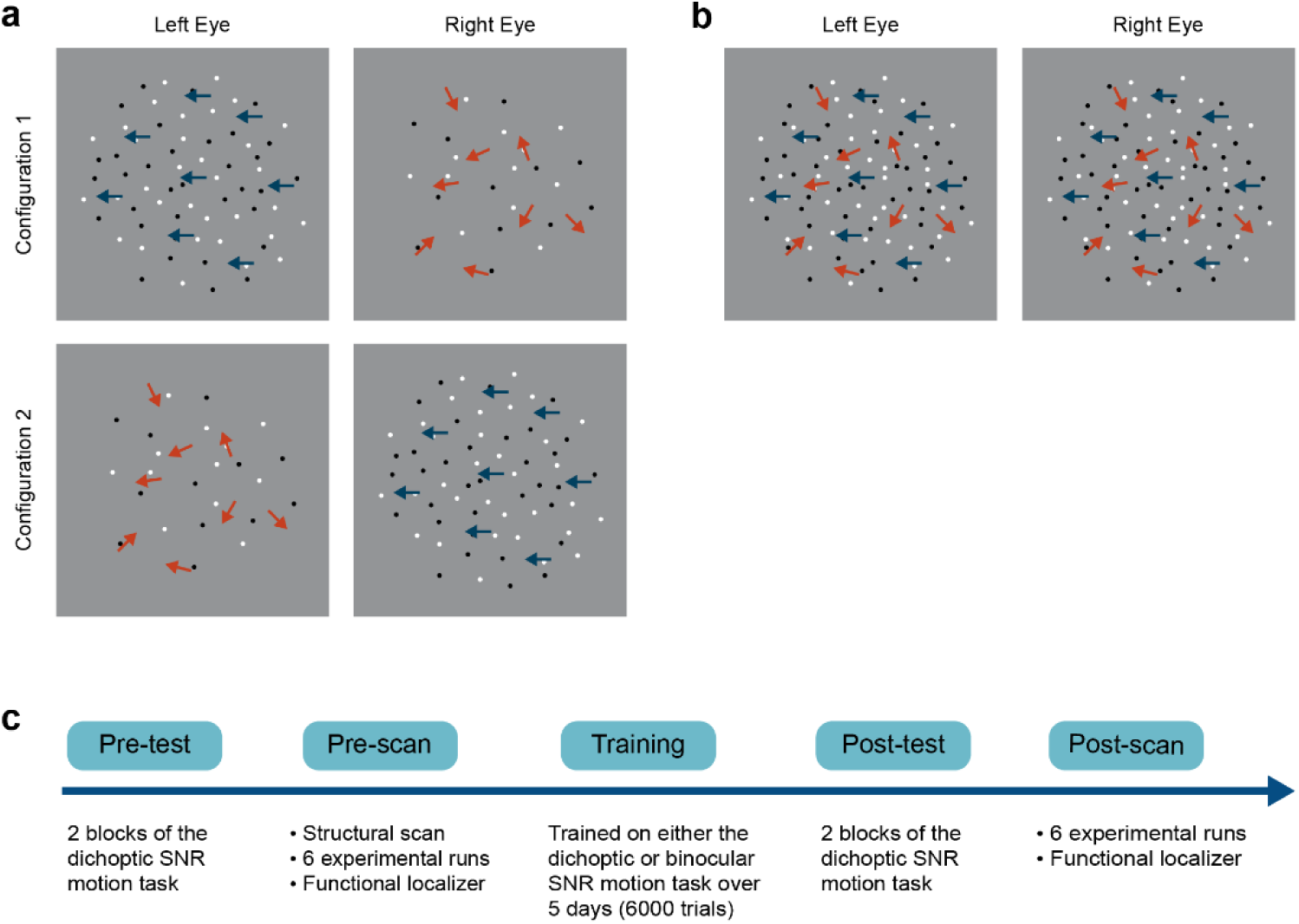
Schematics of the (a) dichoptic and (b) binocular variant of a signal-in-noise (SNR) motion task and (c) the general experimental procedure. For the dichoptic variant, signal and noise dots were presented to different eyes on each trial. Two configurations were used such that we presented signal dots to either the left (configuration 1) or the right eye (configuration 2) on each trial. For the binocular variant, signal and noise dots were presented to both eyes on each trial.

**Figure 2.**
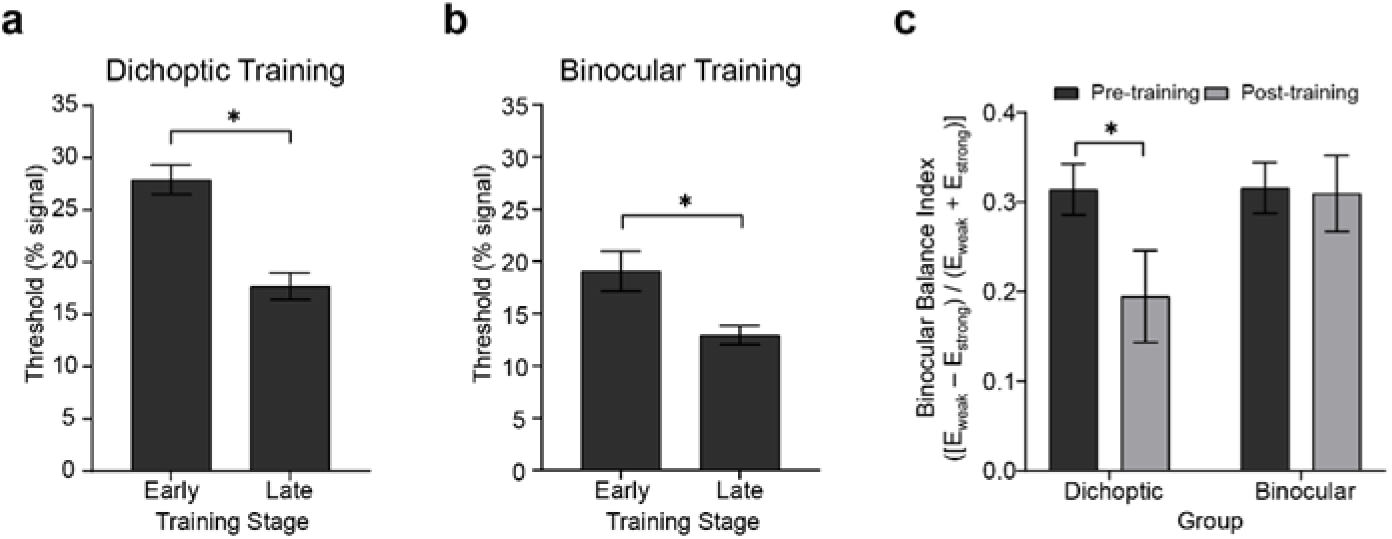
Behavioural results showing the degree of learning and changes in sensory eye dominance. Early and late training thresholds for the (a) dichoptic (N = 24) and (b) binocular (N = 25) training groups. Thresholds for early and late training were derived from averaging the first and the last three training blocks, respectively. (c) Sensory eye dominance in the pre- and post-test for the two training groups as indexed by the binocular balance index derived from the dichoptic signal-in-noise motion test task. An index of zero represents no dominance. Error bars represent ±1 SEM. * p < .05.

We performed two analyses to compare learning effectiveness between training groups. First, we fit individual observer training data with a single-parameter logarithmic function: 𝑏 = 𝑘 ln(𝑎), where 𝑎 and 𝑏 represent the training block and motion threshold, respectively, to determine estimates of the learning rate (𝑘). The learning rate parameter did not significantly differ between the two training groups (t_(47)_ = −1.742, p = 0.090). Second, we compared threshold changes for the respective training tasks (computed as [(thresh_early training_ ― thresh_late training_) / thresh_early training_]). The analysis indicated no significant differences between the two groups in terms of absolute threshold changes (t_(47)_ = 0.970, p = 0.337).

Next, we examined changes in *sensory eye dominance* attained after the 5-day visual training protocol. As in our previous behavioural work (Kam and Chang, 2021), we quantified sensory eye dominance by deriving a binocular balance index from the dichoptic signal-in-noise motion task. The binocular balance index was computed as (*E*_weak_ − *E*_strong_) / (*E*_weak_ + *E*_strong_), where *E*_weak_ represented the threshold obtained when the signal dots were presented to the non-dominant eye, and *E*_strong_ represented the thresholds obtained when the signal dots were presented to the dominant eye. An index of zero represented no dominance, and the more the index deviated from zero, the stronger the dominance. For each participant, the dominant eye was identified in the pre-test, therefore yielding a positive binocular balance index in the pre-test. An index closer to zero would indicate a reduction in dominance after training, while any negative value would represent a change of the dominant eye.

Binocular balance indices were analyzed using a 2 (Group – dichoptic/binocular) × 2 (Time – before/after training) repeated-measures ANOVA that indicated a significant group × time interaction (F_(1, 47)_ = 4.13, p = 0.048, n^2^ = 0.081; Figure 2c). Follow-up paired t-tests revealed that only the group who received dichoptic training demonstrated a reduction in the dichoptic motion task-derived binocular balance indices post-training (t_(23)_ = 2.66, p = 0.014). No change in binocular balance indices was observed in the binocular training group (t_(24)_ = 0.185, p = 0.855). Notably, the baseline (pre-test) binocular balance index was not significantly different between the two groups (t_(47)_ = − 0.044, p = 0.965). Further, training performance of both groups reached asymptotic levels halfway through the training protocol (∼25th block; Figure 3), and learning effectiveness (*k*, above) was comparable between the two groups. Therefore, it is unlikely for the learning-driven changes in sensory eye dominance observed here for the dichoptic-training group to be attributed to dissimilar baseline performances between the two groups or more efficient training for the dichoptically trained group.

**Figure 3.**
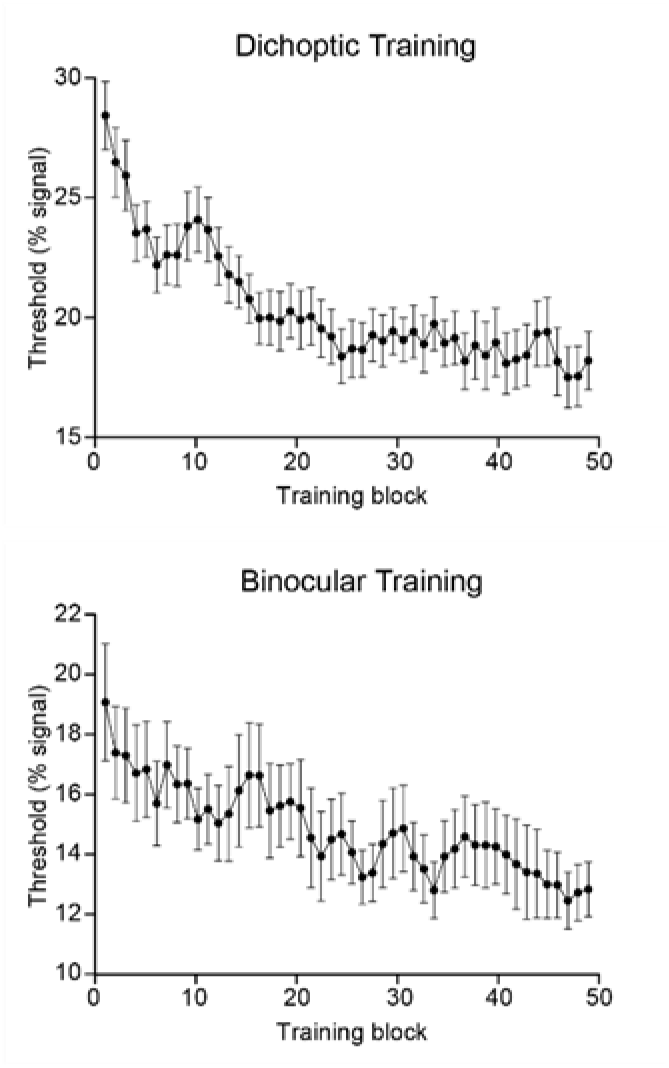
Group-averaged training data presented independently for each training group. Each point represents a three-block moving average. Error bars represent ±1 SEM.

Next, we examined the test-retest reliability of the dichoptic signal-in-noise motion test in quantifying sensory eye dominance by correlating the binocular balance indices obtained from the two test runs, independently for pre- and post-training data using a Pearson’s correlation. The analysis indicated a significant positive correlation between the two test runs (pre-training, r_(47)_ = 0.584, p <.001; post-training, r_(47)_ = 0.688, p < .001).

### fMRI

#### General Linear Model (GLM)

Turning now to the fMRI data, we first evaluated univariate responses in each ROI. For each ROI, GLM beta weights (percent signal change) corresponding to the two stimulus configurations (signal dots presented to the dominant eye and signal dots presented to the non-dominant eye) were extracted for each hemisphere and subsequently entered into a 2 (Group – dichoptic/binocular) × 2 (Time – before/after training) × 2 (Stimulus Configurations – signal dots presented to the dominant eye/signal dots presented to the non-dominant eye) × 2 (Hemisphere – left/right) × 3 (ROI – LGN/V1/hMT+) repeated-measures ANOVA (Figure 4a & b). The analysis revealed a significant main effect of ROI (F_(1.48, 69.79)_ = 21.016, p < .001, n^2^ = 0.309) and hemisphere (F_(1, 47)_ = 7.336, p = 0.009, n^2^ = 0.135). Follow-up Bonferroni-corrected comparisons of the beta weights indicated that the LGN (t_(48)_ = 3.34, p = 0.002) and hMT+ (t_(48)_ = 5.74, p < 0.001) showed significantly higher responses than V1. In general, signals were stronger in the right hemisphere (M = 0.279, SD = 0.407) than the left hemisphere (M = 0.184, SD = 0.448). Univariate responses did not differ between groups (F_(1, 47)_ = 0.903, p = 0.347, n^2^ = 0.019), time (F_(1, 47)_ = 0.179, p = 0.674, n^2^ = 0.004), and conditions (F_(1, 47)_ = 0.051, p = 0.823, n^2^ = 0.001). There were no significant interactions. Although the univariate results indicated stronger activations in the right hemisphere compared to the left hemisphere, we elected to concatenate data from the two hemispheres for the subsequent multivariate pattern analyses owing to the fact that univariate amplitudes are removed in multivariate computations.

**Figure 4.**
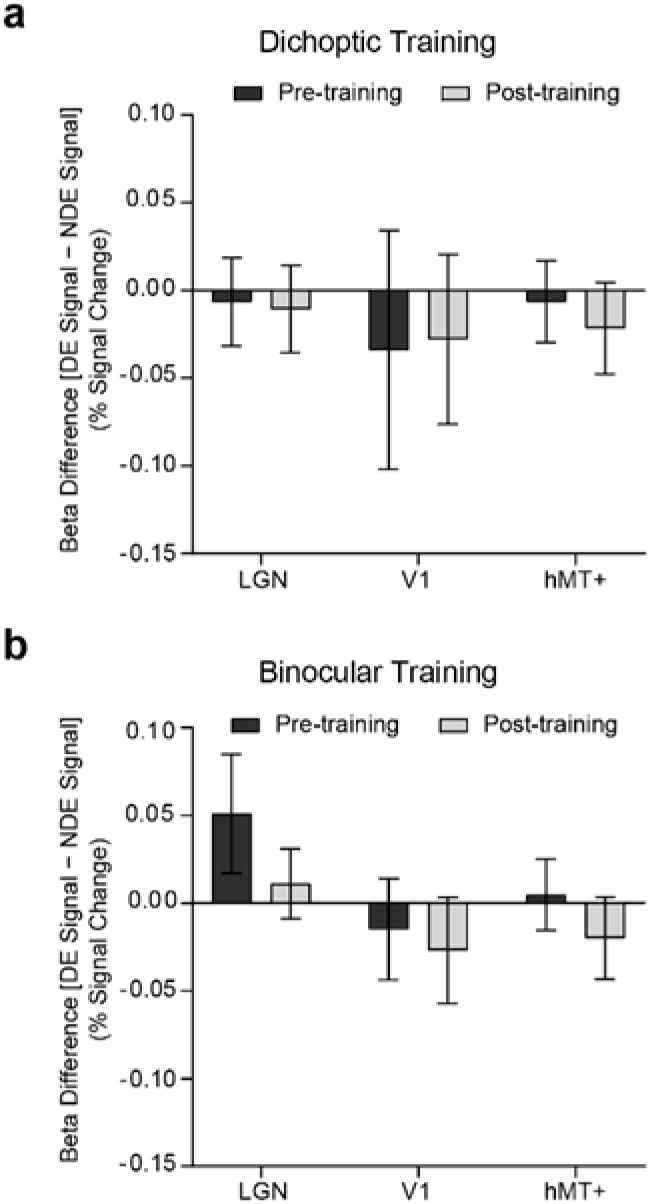
Differences in GLM beta weights [signals presented to the dominant eye (DE signal) – signals presented to the non-dominant eye (NDE signal)] before and after training, presented independently for the (a) dichoptic and (b) binocular training group. Error bars represent ±1 SEM.

#### Multivariate Pattern Analysis (MVPA)

We performed ROI-based multivariate pattern analyses (MVPA) to contrast multivoxel response patterns for the two stimulus configurations [signal dots presented to the dominant eye versus signal dots presented to the non-dominant eye], before and after training (Figure 5). For each (pre and post) dataset and each ROI, classification accuracies were tested against the permutated baseline of 0.50, using t-tests while holding false discovery rate (q < 0.05).

**Figure 5.**
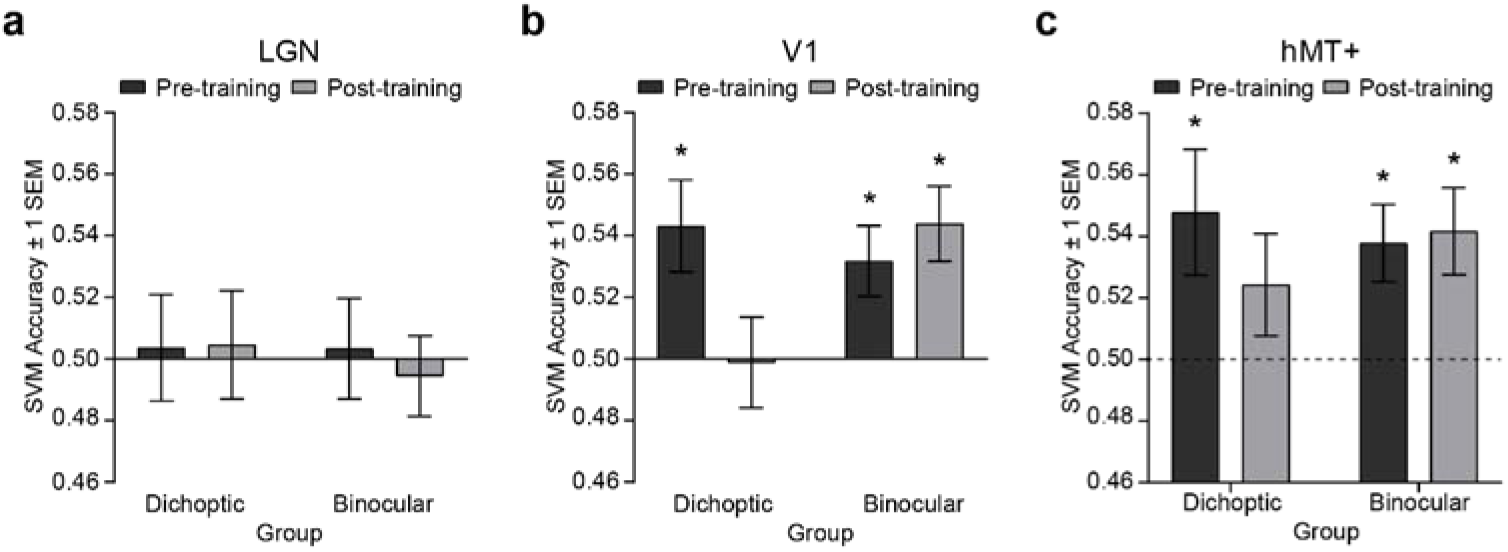
SVM classification accuracies for discriminating fMRI patterned responses between the two stimulus configurations, i.e. [(signal dots presented to the dominant eye) vs (signal dots presented to the non-dominant eye)] before and after training, presented independently for the (a) LGN, (b) V1 and (c) hMT+. Asterisks denote above-baseline (0.5) accuracies. Error bars represent ±1 SEM.

Before perceptual training, accuracies for classifying between signals presented to the dominant eye versus signal presented to the non-dominant eye were significantly above-baseline in V1 (dichoptic, t_(23)_ = 2.89, p = 0.008; binocular, t_(24)_ = 2.773, p = 0.011) and hMT+ (dichoptic, t_(23)_ = 2.336, p =0.029; binocular, t_(24)_ = 3.007, p = 0.006), but not in the LGN (dichoptic, t_(23)_ = −0.204, p = 0.968; binocular, t_(24)_ = −0.586, p = 0.564) *for both groups*. After perceptual training, classification accuracy of the LGN remained at chance level for both groups (dichoptic, t_(23)_ = 0.263 p = 0.795; binocular, t_(24)_ = −0.429, p = 0.674) but classification accuracy of V1 and hMT+ diverged between the two training groups. For the dichoptic training group, SVM accuracies post-training were no longer above-baseline in V1 (t_(23)_ = −0.079, p = 0.938) and hMT+ (t_(23)_ = 1.462, p = 0.157). By contrast, for the binocular training group, SVM accuracies in these two regions remained significantly above-baseline (V1, t_(24)_ = 3.604, p = 0.001; hMT+, t_(24)_ = 2.96, p = 0.007). To further compare pre-and post-training SVM accuracies, we conducted separate, corrected 2 (Group – dichoptic/binocular training) x 2 (Time: pre/post) repeated-measures ANOVAs for each ROI. The ANOVAs revealed a significant Group x Time interaction for V1 only (F_(1, 47)_ = 6.24, p = 0.016, n^2^ = 0.117). Follow-up comparisons indicated that SVM accuracies in V1 were reduced post versus pre-training, specifically for those who received dichoptic perceptual training (t_(23)_ = 3.24, p = 0.004). No changes in SVM accuracies post vs pre-training were observed in the binocular training group (t_(24)_ = −0.68, p = 0.503). Overall, the MVPA results indicated that differences in neural responses between the two stimulus configurations within V1 and hMT+ that existed before training were no longer evident after *dichoptic* visual training. In particular, neural responses of V1 changed significantly following *dichoptic* visual training.

Notably, the observed changes in patterned responses after dichoptic perceptual training could not be simply due to the different signal strengths (i.e. signal-to-noise ratio) of the stimuli, or differences in task difficulty, between pre versus post-fMRI scans. The stimuli used for the experimental runs in the bore were individually tailored based on the thresholds obtained from a behavioural run done inside the bore before (pre and post) image acquisition. While the thresholds obtained at the post-scan were lower for both groups, the differences in thresholds between pre and post-scans (thresh_pre_ – thresh_post_) were not significantly different between the two groups (t_(47)_ = −0.247, p = 0.806). That is, stimulus-level changes were comparable between the two groups, yet changes in the fMRI patterned responses were observed only in those who were *dichoptically* trained. We further verified that our results could not be due to simple differences in task difficulty. To do so, we entered the accuracies of behavioural responses obtained in-bore during the experimental runs into a 2 (Time) x 2 (Group) repeated-measures ANOVA. The analysis revealed no significant effects of group, time, nor interaction, indicating that both groups performed equally well at post-scan versus the pre-scan, even though the signal strength of the stimuli was lower than that of pre-scan. Task performance was also comparable between groups. It is very unlikely, therefore, that the changes in patterned responses after dichoptic perceptual training merely reflect the differences in stimulus difficulty between pre versus post-scans.

Moreover, we verified that the SVM accuracies obtained here were not simply driven by the differences in thresholds between the two stimulus configurations. To test for this, we correlated threshold differences between the two stimulus configurations used in the bore and the SVM accuracies in V1 and hMT+, independently for the pre-training and post-training data. The correlational analyses indicated no significant relationship between the in-bore threshold differences and SVM accuracies in V1 (pre-training, r_(47)_ = 0.022, p = 0.88, post-training, r_(47)_ = −0.121, p = 0.407) and hMT+ (pre-training, r_(47)_ = 0.07, p = 0.633; post-training, r_(47)_ = −0.031, p = 0.831).

#### Brain-behaviour Correlations

As noted above, the patterned responses of V1 and hMT+ were no longer distinguishable between the two stimulus configurations after dichoptic (but not binocular) visual training. In order to better understand the functional relevance of these two brain regions for sensory eye dominance and learning-related changes, we performed an additional set of correlational analyses comparing binocular balance indices and SVM accuracies. Any data points with a Cook’s distance larger than 4/n (where n is the total number of data points) were excluded from the correlational analyses.

We took the absolute value of the post-training binocular balance index such that both pre-and post-training binocular balance indices represented the *degree* of eye dominance. This was done for two reasons: the visual cortex is not concerned with which eye is dominant but rather with the degree of dominance (Muckli et al., 2006). More importantly, SVM accuracy is a non-signed measure that captures only the discriminability of patterned responses between the two stimulus configurations. The dominant eye (left or right) or the change of it is not reflected in SVM accuracy. Therefore, considering only the *degree* of eye dominance (regardless of whether a signage change happened after training) would be more appropriate here. We calculated the correlation coefficient between individual subjects’ SVM accuracies within V1 and hMT+ and the degree of eye dominance, independently for pre- and post-training data (Figure 6). We found that SVM accuracies in V1 (dichoptic, r_(19)_ = 0.473, p = 0.015; binocular, r_(21)_ = 0.437, p = 0.019) and hMT+ (dichoptic, r_(19)_ = 0.421, p = 0.029; binocular, r_(20)_ = 0.513, p = 0.007) correlated positively to binocular balance index *for both groups*, but only before training. In other words, individuals with stronger eye dominance tended to show higher pattern-discriminability for the two stimulus configurations in these two regions. After training, the positive correlation between SVM accuracies and the degree of eye dominance disappeared for the dichoptic training group (V1, r_(19)_ = −0.219, p = 0.170; hMT+, r_(19)_ = −0.168, p = 0.233) but remained for the binocular training group (V1, r_(21)_ = 0.478, p = 0.011; hMT+, r_(20)_ = 0.403, p = 0.032). The correlation between post-training hMT+ SVM accuracies and degree of dominance (post-training) for the binocular training group did not survive statistical correction (q < 0.05).

**Figure 6.**
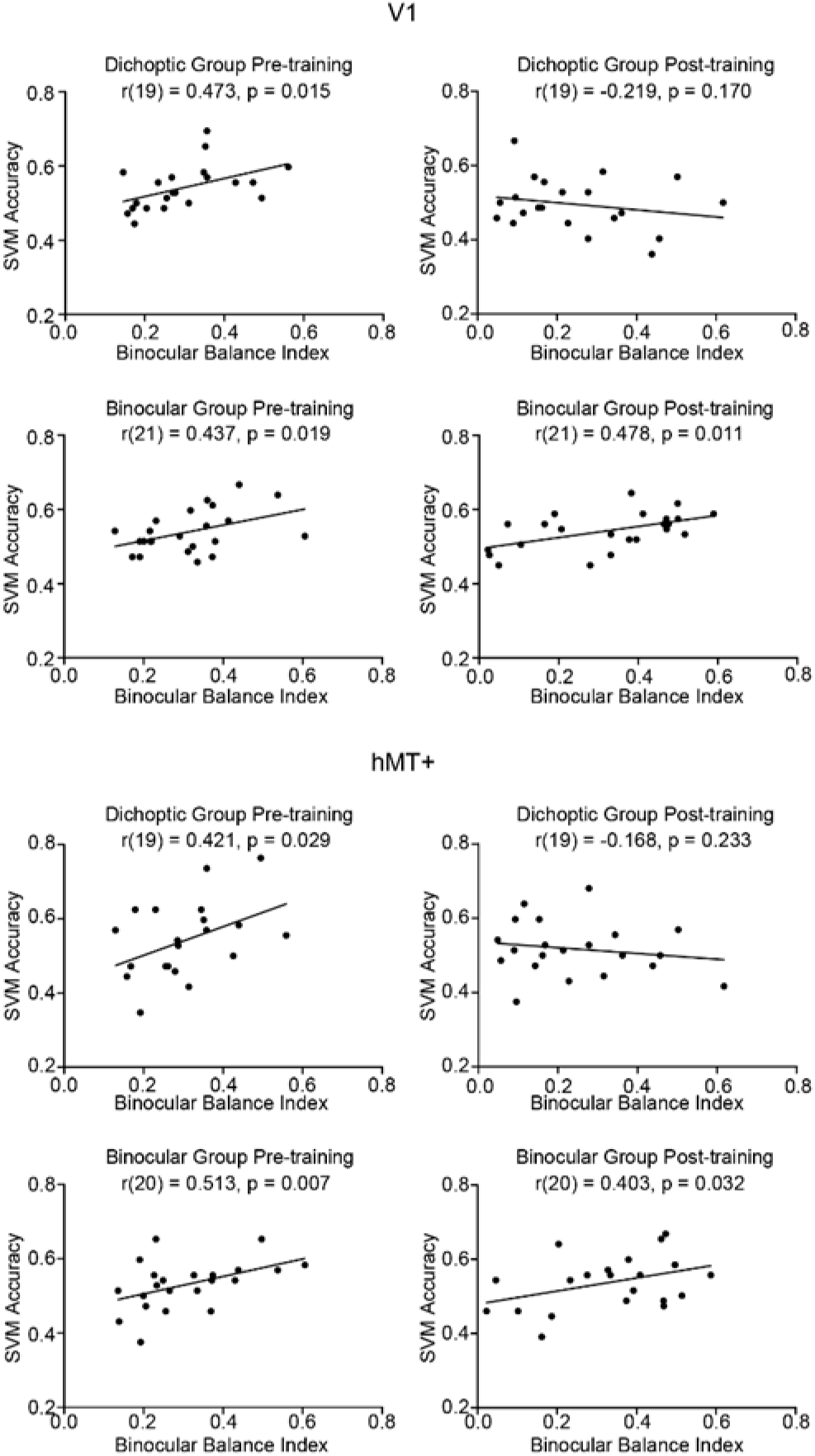
Correlations between SVM accuracies and binocular balance index in V1 and hMT+ before and after training, presented independently for the two training groups.

Finally, we conducted a separate set of correlational analyses on the post-training data considering only those observers whose post-training binocular balance indices remained unchanged in signage (i.e. no change of the dominant eye). The analyses indicated the same pattern of results as that described above, i.e. SVM accuracy remained positively correlated with the degree of eye dominance only for the binocular training group (V1, r_(19)_ = 0.443, p = 0.02; hMT+, r_(19)_ = 0.366, p = 0.05), but not for the dichoptic training group (V1, r_(11)_ = −0.134, p = 0.317; hMT+, r_(11)_ = −0.47, p = 0.439).

## Discussion

We examined the neural mechanisms underlying sensory eye dominance plasticity in binocularly normal adults by measuring changes in fMRI responses before and after a five-day visual training protocol. Specifically, we contrasted training-related changes following dichoptic visual training with changes following training on a binocular variant of the same signal-in-noise motion task. We asked whether learning-driven changes in sensory eye dominance were accompanied by alterations in fMRI responses within the LGN, early retinotopic area (V1), and/or stimulus-specific higher-order extrastriate area (hMT+). Whereas both training groups showed improvements on their dedicated training task, shifts in sensory eye dominance were observed only for those dichoptically-trained. The univariate fMRI responses of the three regions of interest showed relatively homogenous activation across stimulus configurations and were largely comparable before and after training. However, results from the pattern level analysis were more revealing. Before training, patterned responses of V1 and hMT+ in both groups were distinguishable between the two stimulus configurations (i.e., whether signal was presented to the strong or weak eye). Interestingly, after dichoptic (but not binocular) visual training, responses of V1 and hMT+ no longer predicted sensory eye dominance.

We consider first, our behavioural findings. In accordance with our previous work, we observed shifts in sensory eye dominance following dichoptic visual training. Instead of comparing the learning effects with a no-training group as we did previously (Kam and Chang, 2021), we included a binocular variant of the same signal-in-noise motion training in the current design. This was done to make the demands on the two groups more comparable, and to verify that any changes in sensory eye dominance after training could not be due to generic motion training. While both variations of signal-in-noise motion training improved performance on the trained task, only the dichoptic training group exhibited shifts in sensory eye dominance following training. The data thus rule out the possibility that learning-driven *eye balance* improvements are attributed to enhancements in motion perception, signal-in-noise extraction, or additional rule-based cognitive enhancements induced by perceptual training, as put forth by previous work involving adults with amblyopia (Zhang et al., 2014).

We next turn to our fMRI data. Results from the GLM indicated greater overall response amplitudes in the right versus the left hemisphere. This falls in line with previous lesion (Delis et al., 1986; Robertson et al., 1988) and fMRI studies (Han et al., 2002; Martinez et al., 1997) that have shown the right hemisphere to be dominant for processing information that requires global integration of visual stimuli, as would be required for signal integration in our tasks. Further, past behavioural work testing performance on a signal-in-noise motion discrimination task in different visual hemifields reported slightly better task performance when the stimuli were presented to the left versus the right visual field, suggesting a modest right hemisphere dominance for motion signal-in-noise processing in visually healthy individuals (Bosworth and Dobkins, 1999).

Next, we consider the MVPA results. Critically, our multivariate results revealed that V1 and hMT+ are implicated in sensory eye dominance, with the relevance of responses of these two regions to eye dominance disappearing after dichoptic perceptual training. Specifically, before perceptual training, responses of V1 and hMT+ appeared to vary when the signals were presented to different eyes, with the differences in responses escalating with the strength of eye dominance. After dichoptic (but not binocular) visual training, the fMRI patterned responses in V1 and hMT+ no longer predicted sensory eye dominance. That is, dichoptic perceptual training drives not only the rebalancing of activity in the primary visual cortex, but also of activity in stimulus-(motion)-relevant extrastriate cortex.

V1 is a particularly interesting region to consider in this work, not because of its role as the primary cortico-geniculate receiving area, but because it is where we find the emergence of binocularity. While the MVPA results appear to suggest that V1 reweighs two eyes’ input after dichoptic perceptual training such that it considers the data from the two eyes in a more balanced way, our results do not provide further information about the exact mechanisms or layer-specific changes in V1. Based on the prevailing understanding of the excitatory and inhibitory circuits involved in binocular combination (e.g. Meese et al., 2006), one of the possible mechanisms underlying the observed effects relates to interocular suppression. Inhibitory interaction between the eyes is one of the mechanisms that has been linked to sensory eye dominance (Huang et al., 2010; Sengpiel et al., 1994), and it is thought to be mediated by the inhibitory gamma-aminobutyric acid (GABA) circuit, as revealed by animal studies (Harauzov et al., 2010; Sengpiel, 2005; Sengpiel et al., 2006). Recently, a study involving normal-sighted human observers found that individuals with stronger eye dominance tended to have a greater interocular difference in GABAergic inhibition in V1, suggesting a different strength of inhibitory influence of one eye over the other eye during active viewing (Ip et al., 2021). The disappearance of the relevance of V1 responses to eye dominance observed in the current study may thus reflect a rebalancing of interocular suppression. Indeed, it has been reported that resting GABA concentration in V1 can be reduced after short-term monocular deprivation, with the degree of reduction correlating with the changes in eye dominance (Lunghi et al., 2015b). Although monocular deprivation and perceptual training may alter sensory eye dominance through different mechanisms, there is reason to speculate that dichoptic perceptual training may act to balance interocular inhibition in V1, presumably by weakening inhibition from the dominant eye, and/or strengthening inhibition from the non-dominant eye.

From the early classic work, it is well documented that information from the two eyes remains largely segregated in the input layer 4 (Blasdel and Fitzpatrick, 1984; Hubel and Wiesel, 1968b) and only converges at the layer above (Hubel and Wiesel, 1968b). Despite the segregation, a recent study reported (both faciliatory and inhibitory) interocular interactions between monocular neurons in the input layer, before the point where monocular signals integrate (Dougherty et al., 2019a). It is possible, therefore, that the resultant rebalancing effects occur either at the monocular neurons in the input layer before excitatory binocular summation or at the binocular neurons in the upper layers after binocular combination. In this vein then, future work involving ultra-high-field, high-spatial-resolution fMRI may be useful to arrive at learning-driven changes at the laminar level (Huber et al., 2015; Kashyap et al., 2018; Olman et al., 2012).

Unlike data from a previous study indicating no changes in hMT+ after 2 hours of monocular deprivation (Binda et al., 2018), we found that the effects of dichoptic training using a motion stimulus extend beyond V1, and are also observable in training-feature-specific higher-order visual mechanisms. Nevertheless, it remains unclear whether such effects involve intrinsic hMT+ plasticity or are simply fed forward from the primary processing stages. Neurophysiological work indicate that the macaque middle temporal area (MT), receives substantial direct input from the primary visual cortex (Born and Bradley, 2005; Maunsell and Van Essen, 1983). Thus, if the information from the two eyes is reweighed in V1 after dichoptic perceptual training, it would not be surprising for hMT+ to exhibit corresponding (inherited) responses. However, there is evidence that MT cells also receive monocular projections directly from the koniocellular layers of the LGN (Nassi and Callaway, 2006; Sincich et al., 2004; Warner et al., 2010).

Neuroimaging studies imaging effective connectivity (Gaglianese et al., 2012) and tractography (Lanyon et al., 2009) have also demonstrated direct connections between the LGN and hMT+ in the human brain. Although the LGN-hMT+ pathway is less prominent than the geniculo-striate route (Gaglianese et al., 2012), its existence leads to the speculation that the learning-related changes in hMT+ observed in this study may not be solely due to the balanced feedforward input from V1 but also reflect an intrinsic signal reweighting in hMT+ and/or carryover changes that brought about from the LGN. Our data, however, do not support the latter mechanism.

At first glance, our behavioural data appear to be in good consensus with the MVPA results (i.e. reduced behavioural binocular balance index accompanied by reduced pattern-discriminability in V1 and hMT+ after dichoptic perceptual training). Upon close examination, one may notice that while the binocular balance index of the dichoptic training group improved (decreased) with training, it did not drop to zero; however, the residual eye dominance is not reflected in the post-training fMRI data. This raises an intriguing question of where the residual eye dominance is represented. A possible answer for this might be related to something we didn’t image here: white matter. Microstructural abnormalities (i.e. reduced structural integrity and greater diffusivity) in the optic radiations have been reported in adults with amblyopia (Allen et al., 2015; Duan et al., 2015). Similarly, the aforementioned diffusion-weighted imaging work also found white matter diffusivity in the optic radiations to predict the degree of sensory eye dominance in visually-normal adults (Chan and Chang, 2022). Therefore, it may be the case that sensory eye dominance is reflected in both grey and white matter. Any residual eye dominance then, could originate from white matter structural differences that could not be targeted by dichoptic training. Evidence of visual-training-induced microscopic changes within the white matter is scarce. However, data from other domains, such as the motor (Scholz et al., 2009) and cognitive (Mackey et al., 2012; Schlegel et al., 2012; Takeuchi et al., 2010) domains, seem to suggest that longer-term training (at least six weeks) is required to drive changes in the white matter properties. For that reason, we speculate that the white matter remains to play a part in eye dominance after training.

To further complicate the story, it is important to consider the fact that LGN receives about 90% of its input from sites other than the retina (Sherman and Guillery, 2002), including a majority of the input originating from V1 (Van Horn et al., 2000). In principle, then, changes in V1 can in turn feedback to influence LGN responses. Our data provided no indications of learning-related changes in the LGN, however. Compared with the visual cortex, evidence of sensory eye dominance plasticity in human subcortical regions is considerably more limited. To our knowledge, the only study to date that has examined eye balance plasticity in the human LGN also failed to detect any changes in BOLD activity after short-term monocular deprivation (Kurzawski et al., 2022). The lack of LGN-related-effects here, and in work of others could reflect a true absence of a role for this region in governing eye balance. Alternatively, it’s important to consider several challenges that the LGN poses:

First, the LGN is much smaller in size (and hence weaker signal-to-noise ratio) as compared to V1 and hMT+. As most of our participants had relatively balanced eyes (even before training), response amplitude differences between stimulus configurations or after learning in the LGN may be more difficult to detect under these circumstances. Another possible explanation for our results is that dichoptic perceptual training may drive changes in subcortical and cortical visual processing via mechanisms that are very different in nature. The LGN and the visual cortex (V1 and hMT+) are fundamentally different in a way that most of the cells in the LGN are monocular (Casagrande and Boyd, 1996), while those in V1 (Hubel and Wiesel, 1968b) and hMT+ (Maunsell and Van Essen, 1983) are mainly binocular. Also, binocular modulations in the LGN are predominately inhibitory or suppressive (Dougherty et al., 2021), but those in V1 can be facilitatory (Dougherty et al., 2019a). Indeed, various rodent studies have indicated that the characteristics of changes observed in the LGN and visual cortex following monocular deprivation are very different. For instance, a study in which miniature excitatory postsynaptic currents (mEPSC) in adult mice were recorded showed that after monocular deprivation, the mEPSC amplitude of deprived-eye neurons increased in both the LGN and the visual cortex (Krahe and Guido, 2011). However, the increased mEPSC amplitude in the LGN was also accompanied by an increase in frequency (Desai et al., 2002), suggesting that experience-dependent plasticity in the LGN and visual cortex could be mediated by different presynaptic and/or postsynaptic mechanisms. It is also worth noting that many of these rodent studies that have detected eye-specific response changes in the LGN mainly used invasive neurophysiological techniques that tracked individual neurons (Desai et al., 2002; Jaepel et al., 2017; Krahe and Guido, 2011), in contrast to the non-invasive but indirect method of fMRI here. It is therefore possible that there are changes in the LGN driven by dichoptic perceptual training here that are not well reflected by BOLD activity.

## Conclusions

Combining behavioural training and fMRI paradigms, we show for the first time how dichoptic perceptual training drives sensory eye dominance plasticity in the visual cortex. Our data suggest that visual training using dichoptic presentation of signal-in-noise motion stimuli leads to changes in sensory eye dominance by potentially driving a reweighting of data from the two eyes in both the primary and task-related extrastriate visual areas. Our findings establish a foundational basis for future work that seeks to better understand training-related improvements at cortex – for example, the interplay between cortical inhibition and excitation. Future research could also explore the generality of training-induced improvements in sensory eye dominance by testing alternative measures of sensory eye dominance, such as a binocular rivalry test.

Reaching a better understanding of binocular plasticity and the neural underpinnings of eye dominance is essential for developing effective rehabilitative paradigms for the visually impaired.

## Materials and Methods

### Participants

Fifty visually normal observers participated in this study (mean age of 22.2 years; SD 3.4 years; 27 males). All had normal or corrected-to-normal visual acuity as screened with the LogMAR chart (20/20) and normal binocular fusion as screened with the Worth-4-dots test. The Worth-4-dots test was performed at 33cm from the observers, who were shown four dots of light arranged in a diamond configuration (one red dot, two green dots, and one white dot). The observers were required to report the number and colour of dots through red/green anaglyph glasses. Normal binocular fusion was indicated by a report of four dots (one red, two green and one mixed colour). All the participants were right-handed. They provided written informed consent in line with the ethical review and approval by the Human Research Ethics Committee (HREC), The University of Hong Kong. Participants were randomly assigned to two training groups — receiving training on either a dichoptic (N = 25; mean age of 23.6 years; SD 3.7 years; 13 males) or binocular (N = 25; mean age of 22.3 years; SD 3.3 years; 14 males) variant of the signal-in-noise motion task. One observer from the dichoptic visual training group was excluded from the final analysis due to extensive head movements during the fMRI scan. The sample size was determined based upon statistical power analysis, using the effect size reported in a previous study that employed the same dichoptic training task (Kam and Chang, 2021), with the aim of attaining a minimum of 80% power to detect learning-related changes.

### General Procedure

A schematic of the experimental procedure is presented in Figure 1c. Both groups completed pre-training and post-training laboratory tests and fMRI scans. Participants were tested on the dichoptic signal-in-noise (SNR) motion task during the pre- and post-tests. They performed the same task at the pre- and post-scans, during which BOLD signals were measured concurrently. The two groups received training on either the dichoptic or binocular signal-in-noise motion task over five consecutive days (6000 trials in total). Each training session lasted 60 minutes. The post-test was done immediately following the last training session (i.e., after the last training block), while the post-scan was completed the day immediately following the last training session.

### In-laboratory Testing and Training

#### Apparatus

Stimuli were generated using custom software written in MATLAB, with extensions from Psychtoolbox (Brainard, 1997; Pelli, 1997). Dichoptic presentation of the stimuli was achieved via a shutter-presentation setup that consisted of an ASUS 3D-vision-ready LCD display (resolution: 1920 x 1080 pixels; refresh rate: 120 Hz) paired with NVIDIA 3D Vision 2 shutter glasses. To ensure complete segregation of the two eyes’ signals, we presented different geometric test patterns independently to each eye at the beginning of each session. The observers were instructed to view the geometric test patterns alternatively with their left and right eyes and to report what they observed. No incidents of crosstalk between eyes were reported. The stimuli were viewed at a distance of 50 cm, which was maintained by a chin-rest.

#### Stimuli and Tasks

Stimuli were presented against a uniform gray background and surrounded by a binocularly presented grid-like frame of white and black squares (each 1.5 deg in size). This grid served to promote binocular fusion by providing an unambiguous background reference. The dichoptic signal-in-noise motion stimuli used in laboratory testing and training were identical to those used in our previous behavioural work (Kam and Chang, 2021). The stimuli consisted of 60 black and 60 white non-overlapping dots presented at 100% contrast. Each dot had a size of 0.2 degrees and moved within a central aperture 9 degrees in diameter (dot density of 1.48 dots/deg^2^) with a velocity of 2 deg/s. The dots did not have a limited lifetime. At the beginning of each trial, the position of each dot was randomly assigned within the aperture. Dots that had an impending collision or were to move outside the aperture on the next update (frame) were redrawn to a random position.

The detectability of motion direction depended on the signal-to-noise ratio, which varied from 0% to 100%. At 100% signal, all the dots moved coherently in either the left or right direction, while at 0% signal, all dots moved in random directions within the aperture. On each trial, we presented signal dots and noise dots dichoptically (i.e., signal and noise dots were presented to different eyes). Observers were asked to make a two-alternative forced-choice judgement of the net motion direction of the dots (either leftward or rightward) by pressing one of two the arrow keys on the keyboard. Task difficulty was manipulated by adjusting the signal-to-noise ratio for each trial using the QUEST staircase procedure, measuring the percentage of signal required to achieve an 82% correctness level. A block of trials consisted of two interleaved staircases of 60 trials, with each presenting signal dots to either the left (Figure 1a, configuration 1) or the right eye (Figure 1a, configuration 2). The two staircases were interleaved such that the eye of origin for signals and noise could not be determined on each trial once the two eyes’ images were fused. Stimuli were presented for 500 ms, followed by a 1000 ms response period. Trials were separated by a 300-ms interval. Participants practiced for 20 trials in the pre-test to become familiar with the task and completed two test blocks in both the pre- and post-laboratory tests. Auditory feedback was given during the training (but not the test) sessions.

All the stimulus and task parameters of the binocular variant of the signal-in-noise motion task were identical to those described above for the dichoptic variant, except that signal and noise dots were presented to both eyes on each trial (Figure 1b). The binocular variant was used in training only and was not tested in the pre- and post- test.

### fMRI acquisition, design, analysis

#### Apparatus

Stimuli were presented dichoptically in the magnet via a projector (ProPixx, Vpixx) fit with a circular polarizer set to display 1920 x 1080 pixels at 120 Hz. The stimuli were side-projected to a standing mirror placed at 45° behind the bore. The mirror image was then projected onto a translucent 3D rear projection screen placed at the base of the bore. Observers viewed the stimuli through a 45° tilted coil-mounted mirror in front of the head with passive polarized filters inserted in the custom MR- compatible frames. The frames were selected based on the subjects’ intra-pupillary distance. A second optional corrective lens was provided when needed (for myopia). Prior to each scanning session, we verified the segregation of the left and right eye channels by displaying geometric test patterns independently to each eye. Behavioural responses were collected using an MR-compatible response box.

#### Stimuli and Task

Participants were scanned during completion of the dichoptic signal-in-noise motion task. The in-bore stimuli were the same as those used in the laboratory except for the following differences: First, the stimuli were presented at 70% contrast. Second, the size of the central aperture was set to 10 deg in diameter, with each dot subtending 0.22 degrees and moving with a velocity of 4 deg/s. This change was made to exaggerate the stimulus while minimizing cross-talk in-bore.

#### fMRI acquisition

Imaging data were acquired using a GE SIGNA Premier 3.0T scanner with a phased array 48-channel head coil. For both experimental runs and functional localizers, blood oxygen level-dependent signals were measured with a multiband echo-planar sequence (voxel size = 2 x 2 x 2 mm^3^; TR = 2000 ms; TE = 30 ms; FOV = 240 x 240; flip angle = 90°; 58 slices; multiband factor = 2; 200 volumes). Additionally, a high-resolution T1-weighted image was acquired for each participant (voxel size = 1 x 1 x 1 mm^3^; TR = 7 ms; TE = 2.8 ms; FOV = 256 x 256, flip angle = 8°).

#### Region of Interest (ROI) Localization

V1 and hMT+ were defined using separate functional localizer scans. We identified V1 using standard phase-encoded retinotopic mapping procedures that mapped polar angles with a slowly rotating checkerboard wedge stimulus (Sereno et al., 1995). Each participant completed clockwise and counterclockwise rotating retinotopic localizer scans. hMT+ was localized using a single-run functional localizer and defined as a cluster of contiguous voxels that showed significantly stronger activation to an array of coherently contracting or expanding dots than to an array of stationary dots (Huk et al., 2002). The LGN was anatomically defined as a 3 mm radius spherical ROI centered on the Talairach coordinate of [left: −22, −24, −2; right: 22, −24 −2] (Chang et al., 2016).

The mean Talairach coordinates of V1 and hMT+ as identified in this study, for each hemisphere, are presented in Table 1. The Talairach coordinates of both regions were in good agreement with those reported previously (Aedo-Jury et al., 2020; Gaglianese et al., 2012; Poghosyan and Ioannides, 2007).

**Table 1.** Talairach coordinates (mean ± SD) of V1 and hMT+.

|  | Left |  |  | Right |  |  |
| --- | --- | --- | --- | --- | --- | --- |
|  | X | Y | Z | X | Y | Z |
| **V1** | -8.98±3.82 | -89.90±1.97 | -5.33±4.45 | 11.12±2.69 | -89.42±1.91 | -0.83±4.14 |
| **hMT+** | -41.66± 3.52 | -65.45±3.54 | -0.43±3.45 | 41.36±2.15 | -63.32±3.18 | 0.44±3.28 |

#### Design and Procedure

Before commencing image acquisition, each participant completed one behaviour-only run of the dichoptic signal-in-noise motion task while laying inside the bore to obtain thresholds used for computing individually tailored stimulus test values for the main experimental runs. This strategy allowed us to match task difficulty across participants and conditions. Similar to the in-lab tests, one run consisted of two interleaved staircases of 60 trials, each corresponding to the two stimulus conditions (signal dots presented to the left or right eye). We averaged the test values in the last 30 trials for each condition and defined a range of stimuli values of ±1 SD from this mean value. For each trial, we then sampled the signal-to-noise ratio from this range.

We adopted a block design for the fMRI runs, with each block lasting 16 sec. Each run comprised three block types: two stimulus configuration blocks (signal presented to the left or right eye) and a fixation block. Each stimulus block consisted of 8 trials. On each trial, the stimulus was presented for 500 ms and was followed by a 1500 ms response period, during which the observers were asked to judge the net motion direction of the dots by pressing buttons on the response box. A fixation block contained a white fixation cross 0.8 deg in size that was presented at the center of the screen for 16 sec. The order of the stimulus configurations was randomized and stimulus blocks were interleaved with fixation blocks. Within a particular run, each stimulus block was repeated six times, yielding 48 repetitions of a particular stimulus configuration and a total run time of 6 min and 40 sec. Each participant completed six fMRI acquisition runs in both the pre- and post-scan. A full scan session lasted around 75 minutes.

#### fMRI Data Analysis

Imaging data were processed using BrainVoyager 22.0. The anatomical data of each participant were transformed into Talairach space and used for cortex reconstruction and inflation. For each functional run, the initial two volumes were discarded to eliminate effects of startup magnetization transients in the data. Functional data were preprocessed using slice scan time correction (with cubic-spline interpolation), 3D head motion correction, linear trend removal, and temporal high-pass filtering (three cycles per run). The preprocessed functional images were subsequently aligned to each participant’s anatomical images and transformed into Talairach space (Talairach and Tournoux, 1988).

We examined univariate responses (general linear modal, GLM) and multivariate pattern responses (multivoxel pattern analysis, MVPA). The GLM included regressors for the fixation and the two stimulus conditions (signal dots presented to the dominant eye and signal dots presented to the non-dominant eye) and six motion regressors (three translation parameters and three rotation parameters). The averaged time course signals obtained across all voxels in each ROI were then modelled as a linear combination of the different regressors. The regressor coefficients or beta weights of different stimulus conditions were used for contrasts of the two stimulus conditions.

Unlike GLM, which considers overall responsivity, MVPA considers pattern-level responses and their uniqueness to the stimulus condition(s). MVPA classification analyses were performed using a linear support vector machine (SVM) classifier. Specifically, the fMRI time course signals of all voxels were first converted to z scores and shifted by 4 sec (2 TRs). This shift was introduced to account for the hemodynamic response delay (Serences, 2004). For each ROI, the SVM was trained to classify the patterned responses between the two stimulus configurations: signal dots presented to the dominant eye versus signal dots presented to the non-dominant eye. We adopted a leave-one-run-out cross-validation procedure for the SVM. In each ROI, the functional data of one run was used as the validation dataset, while the remaining runs were used as the training dataset. This analysis was repeated 17 times at all possible fine voxel counts between 10 and 800 voxels (stepping on 50 voxels increment), each computing a classification accuracy at the corresponding voxel count. The final pattern size reported here was determined by the smallest pattern size at which accuracies reached asymptotic levels, corresponding to 400 voxels for our dataset. For each ROI, the mean classification accuracies at 400 voxels were tested against the chance level (0.50), as computed by running 1000 SVMs with shuffled labels.

## Data/code availability

Anonymized behavioural data are available at the data repository of the University of Hong Kong, https://doi.org/10.25442/hku.22662559. The conditions of our ethics approval do not permit the public archiving of raw fMRI data because the raw data files contain subject-identifiable information that are inherent to the data themselves (e.g., skull data), and in the headers. There is no method available to eliminate this identifiable information while retaining the data in their rawest form (i.e., without some degree of preprocessing). The imaging data, however, can be obtained by readers from the corresponding author via written request but require additional approval to be sought from the Human Research Ethics Committee of The University of Hong Kong. Custom code written for the presentation of stimuli and/or analyses of data in this manuscript are available from the corresponding authors upon reasonable request.

## References

Aedo-Jury, F., Cottereau, B.R., Celebrini, S., Séverac Cauquil, A., 2020. Antero-posterior vs. lateral vestibular input processing in human visual cortex. Frontiers in integrative neuroscience 14, 43.

Allen, B., Spiegel, D.P., Thompson, B., Pestilli, F., Rokers, B., 2015. Altered white matter in early visual pathways of humans with amblyopia. Vision research 114, 48–55.

Beckers, G., Hömberg, V., 1992. Cerebral visual motion blindness: transitory akinetopsia induced by transcranial magnetic stimulation of human area V5. Proceedings of the Royal Society of London. Series B: Biological Sciences 249, 173–178.

Binda, P., Kurzawski, J.W., Lunghi, C., Biagi, L., Tosetti, M., Morrone, M.C., 2018. Response to short-term deprivation of the human adult visual cortex measured with 7T BOLD. Elife 7, e40014.

Blasdel, G.G., Fitzpatrick, D., 1984. Physiological organization of layer 4 in macaque striate cortex. Journal of Neuroscience 4, 880–895.

Born, R.T., Bradley, D.C., 2005. Structure and function of visual area MT. Annual review of neuroscience 28, 157.

Bosworth, R.G., Dobkins, K.R., 1999. Left-hemisphere dominance for motion processing in deaf signers. Psychological Science 10, 256–262.

Braddick, O.J., O’Brien, J.M., Wattam-Bell, J., Atkinson, J., Hartley, T., Turner, R., 2001. Brain areas sensitive to coherent visual motion. Perception 30, 61–72.

Brainard, D.H., 1997. The psychophysics toolbox. Spatial vision 10, 433–436. https://doi.org/10.1163/156856897X00357

Casagrande, V.A., Boyd, J.D., 1996. The neural architecture of binocular vision. Eye 10, 153–160.

Chan, A.Y., Chang, D.H., 2022. Neural Correlates of Sensory Eye Dominance in Human Visual White Matter Tracts. Eneuro.

Chang, D.H., Hess, R.F., Mullen, K.T., 2016. Color responses and their adaptation in human superior colliculus and lateral geniculate nucleus. Neuroimage 138, 211– 220.

Conner, I.P., Odom, J.V., Schwartz, T.L., Mendola, J.D., 2007. Monocular activation of V1 and V2 in amblyopic adults measured with functional magnetic resonance imaging. Journal of American Association for Pediatric Ophthalmology and Strabismus 11, 341–350. https://doi.org/10.1016/j.jaapos.2007.01.119

Delis, D.C., Robertson, L.C., Efron, R., 1986. Hemispheric specialization of memory for visual hierarchical stimuli. Neuropsychologia 24, 205–214.

Desai, N.S., Cudmore, R.H., Nelson, S.B., Turrigiano, G.G., 2002. Critical periods for experience-dependent synaptic scaling in visual cortex. Nature neuroscience 5, 783–789.

Ding, J., Klein, S.A., Levi, D.M., 2013. Binocular combination in abnormal binocular vision. Journal of vision 13, 14–14. https://doi.org/10.1167/13.2.14

Dougherty, K., Carlson, B.M., Cox, M.A., Westerberg, J.A., Zinke, W., Schmid, M.C., Martin, P.R., Maier, A., 2021. Binocular Suppression in the Macaque Lateral Geniculate Nucleus Reveals Early Competitive Interactions between the Eyes. eNeuro 8, ENEURO.0364-20.2020. https://doi.org/10.1523/ENEURO.0364-20.2020

Dougherty, K., Cox, M.A., Westerberg, J.A., Maier, A., 2019a. Binocular modulation of monocular V1 neurons. Current Biology 29, 381–391.

Dougherty, K., Schmid, M.C., Maier, A., 2019b. Binocular response modulation in the lateral geniculate nucleus. Journal of Comparative Neurology 527, 522–534.

Duan, Y., Norcia, A.M., Yeatman, J.D., Mezer, A., 2015. The structural properties of major white matter tracts in strabismic amblyopia. Investigative ophthalmology & visual science 56, 5152–5160.

Freeman, R.D., Tsumoto, T., 1983. An electrophysiological comparison of convergent and divergent strabismus in the cat: electrical and visual activation of single cortical cells. Journal of Neurophysiology 49, 238–253.

Gaglianese, A., Costagli, M., Bernardi, G., Ricciardi, E., Pietrini, P., 2012. Evidence of a direct influence between the thalamus and hMT+ independent of V1 in the human brain as measured by fMRI. Neuroimage 60, 1440–1447.

Gaska, A.W.P.J.P., Foote, W., Pollen, D.A., 2000. Striate cortex increases contrast gain of macaque LGN neurons. Visual neuroscience 17, 485494. https://doi.org/10.1017/s0952523800174012

Guillery, R.W., Colonnier, M., 1970. Synaptic patterns in the dorsal lateral geniculate nucleus of the monkey. Zeitschrift für Zellforschung und mikroskopische Anatomie 103, 90–108.

Han, S., Weaver, J.A., Murray, S.O., Kang, X., Yund, E.W., Woods, D.L., 2002. Hemispheric asymmetry in global/local processing: effects of stimulus position and spatial frequency. Neuroimage 17, 1290–1299.

Harauzov, A., Spolidoro, M., DiCristo, G., De Pasquale, R., Cancedda, L., Pizzorusso, T., Viegi, A., Berardi, N., Maffei, L., 2010. Reducing intracortical inhibition in the adult visual cortex promotes ocular dominance plasticity. Journal of Neuroscience 30, 361–371.

Hendrickson, A.E., Wilson, J.R., Ogren, M.P., 1978. The neuroanatomical organization of pathways between the dorsal lateral geniculate nucleus and visual cortex in Old World and New World primates. The Journal of comparative neurology 182, 123–136.

Hess, R.F., Mansouri, B., Thompson, B., 2010. A new binocular approach to the treatment of amblyopia in adults well beyond the critical period of visual development. Restorative neurology and neuroscience 28, 793–802. https://doi.org/10.3233/RNN-2010-0550

Hess, R.F., Thompson, B., Gole, G., Mullen, K.T., 2009. Deficient responses from the lateral geniculate nucleus in humans with amblyopia. European Journal of Neuroscience 29, 1064–1070.

Huang, C.-B., Zhou, J., Zhou, Y., Lu, Z.-L., 2010. Deficient Binocular Combination Reveals Mechanisms of Anisometropic Amblyopia. Journal of Vision 10, 466– 466.

Hubel, D.H., Wiesel, T.N., 1972. Laminar and columnar distribution of geniculo-cortical fibers in the macaque monkey. Journal of Comparative Neurology 146, 421–450.

Hubel, D.H., Wiesel, T.N., 1968a. Receptive fields and functional architecture of monkey striate cortex. The Journal of physiology 195, 215–243. https://doi.org/10.1113/jphysiol.1968.sp008455

Hubel, D.H., Wiesel, T.N., 1968b. Receptive fields and functional architecture of monkey striate cortex. The Journal of physiology 195, 215–243.

Huber, L., Goense, J., Kennerley, A.J., Trampel, R., Guidi, M., Reimer, E., Ivanov, D., Neef, N., Gauthier, C.J., Turner, R., Möller, H.E., 2015. Cortical lamina-dependent blood volume changes in human brain at 7T. NeuroImage 107, 23–33. https://doi.org/10.1016/j.neuroimage.2014.11.046

Huk, A.C., Dougherty, R.F., Heeger, D.J., 2002. Retinotopy and functional subdivision of human areas MT and MST. Journal of Neuroscience 22, 7195–7205.

Ip, I.B., Emir, U.E., Lunghi, C., Parker, A.J., Bridge, H., 2021. GABAergic inhibition in the human visual cortex relates to eye dominance. Scientific reports 11, 1–11.

Jaepel, J., Hübener, M., Bonhoeffer, T., Rose, T., 2017. Lateral geniculate neurons projecting to primary visual cortex show ocular dominance plasticity in adult mice. Nature neuroscience 20, 1708–1714.

Kam, K.Y., Chang, D.H.F., 2021. Dichoptic Perceptual Training and Sensory Eye Dominance Plasticity in Normal Vision. Investigative Ophthalmology & Visual Science 62, 12. https://doi.org/10.1167/iovs.62.7.12

Kashyap, S., Ivanov, D., Havlicek, M., Sengupta, S., Poser, B.A., Uludağ, K., 2018. Resolving laminar activation in human V1 using ultra-high spatial resolution fMRI at 7T. Sci Rep 8, 17063. https://doi.org/10.1038/s41598-018-35333-3

Krahe, T.E., Guido, W., 2011. Homeostatic plasticity in the visual thalamus by monocular deprivation. Journal of Neuroscience 31, 6842–6849.

Kurzawski, J.W., Lunghi, C., Biagi, L., Tosetti, M., Morrone, M.C., Binda, P., 2022. Short-term plasticity in the human visual thalamus. eLife 11, e74565. https://doi.org/10.7554/eLife.74565

Lanyon, L.J., Giaschi, D., Young, S.A., Fitzpatrick, K., Diao, L., Bjornson, B.H., Barton, J.J., 2009. Combined functional MRI and diffusion tensor imaging analysis of visual motion pathways. Journal of Neuro-Ophthalmology 29, 96–103.

Li, J., Lam, C.S., Yu, M., Hess, R.F., Chan, L.Y., Maehara, G., Woo, G.C., Thompson, B., 2010. Quantifying sensory eye dominance in the normal visual system: a new technique and insights into variation across traditional tests. Investigative Ophthalmology & Visual Science 51, 6875–6881. https://doi.org/10.1167/iovs.10-5549

Li, J., Thompson, B., Deng, D., Chan, L.Y., Yu, M., Hess, R.F., 2013. Dichoptic training enables the adult amblyopic brain to learn. Current Biology 23, R308–R309.

Lunghi, C., Berchicci, M., Morrone, M.C., Di Russo, F., 2015a. Short-term monocular deprivation alters early components of visual evoked potentials: Homeostatic plasticity in adult visual cortex. J Physiol 593, 4361–4372. https://doi.org/10.1113/JP270950

Lunghi, C., Emir, U.E., Morrone, M.C., Bridge, H., 2015b. Short-term monocular deprivation alters GABA in the adult human visual cortex. Current Biology 25, 1496–1501.

Mackey, A.P., Whitaker, K.J., Bunge, S.A., 2012. Experience-dependent plasticity in white matter microstructure: reasoning training alters structural connectivity. Frontiers in neuroanatomy 6, 32.

Marrocco, R.T., McClurkin, J.W., 1979. Binocular interaction in the lateral geniculate nucleus of the monkey. Brain research 168, 633–637.

Martinez, A., Moses, P., Frank, L., Buxton, R., Wong, E., Stiles, J., 1997. Hemispneric asymmetries in global and local processing: evidence from fMRI. Neuroreport 8, 1685–1689.

Maunsell, J.H., Van Essen, D.C., 1983. Functional properties of neurons in middle temporal visual area of the macaque monkey. II. Binocular interactions and sensitivity to binocular disparity. Journal of neurophysiology 49, 1148–1167. https://doi.org/10.1152/jn.1983.49.5.1148

Meese, T.S., Georgeson, M.A., Baker, D.H., 2006. Binocular contrast vision at and above threshold. Journal of vision 6, 7–7. https://doi.org/10.1167/6.11.7

Muckli, L., Kieß, S., Tonhausen, N., Singer, W., Goebel, R., Sireteanu, R., 2006. Cerebral correlates of impaired grating perception in individual, psychophysically assessed human amblyopes. Vision research 46, 506–526.

Nassi, J.J., Callaway, E.M., 2006. Multiple circuits relaying primate parallel visual pathways to the middle temporal area. Journal of Neuroscience 26, 12789– 12798.

Olman, C.A., Harel, N., Feinberg, D.A., He, S., Zhang, P., Ugurbil, K., Yacoub, E., 2012. Layer-specific fMRI reflects different neuronal computations at different depths in human V1. PloS one 7, e32536.

Pelli, D.G., 1997. The VideoToolbox software for visual psychophysics: Transforming numbers into movies. Spatial vision 10, 437–442. https://doi.org/10.1163/156856897X00366

Poghosyan, V., Ioannides, A.A., 2007. Precise mapping of early visual responses in space and time. Neuroimage 35, 759–770.

Ramamurthy, M., Blaser, E., 2021. The ups and downs of sensory eye balance: Monocular deprivation has a biphasic effect on interocular dominance. Vision Research 183, 53–60. https://doi.org/10.1016/j.visres.2021.01.010

Robertson, L.C., Lamb, M.R., Knight, R.T., 1988. Effects of lesions of temporal-parietal junction on perceptual and attentional processing in humans. Journal of Neuroscience 8, 3757–3769.

Rodieck, R.W., Dreher, B., 1979. Visual suppression from nondominant eye in the lateral geniculate nucleus: a comparison of cat and monkey. Experimental Brain Research 35, 465–477.

Sanderson, K.J., Bishop, P.O., Darian-Smith, I., 1971. The properties of the binocular receptive fields of lateral geniculate neurons. Experimental Brain Research 13, 178–207. https://doi.org/10.1007/BF00234085

Schlegel, A.A., Rudelson, J.J., Tse, P.U., 2012. White matter structure changes as adults learn a second language. Journal of cognitive neuroscience 24, 1664– 1670.

Scholz, J., Klein, M.C., Behrens, T.E.J., Johansen-Berg, H., 2009. Training induces changes in white-matter architecture. Nat Neurosci 12, 1370–1371. https://doi.org/10.1038/nn.2412

Sengpiel, F., 2005. Intracortical Origins of Interocular Suppression in the Visual Cortex. Journal of Neuroscience 25, 6394–6400. https://doi.org/10.1523/JNEUROSCI.0862-05.2005

Sengpiel, F., Blakemore, C., Harrad, R., 1995. Interocular suppression in the primary visual cortex: a possible neural basis of binocular rivalry. Vision Research 35, 179–195. https://doi.org/10.1016/0042-6989(94)00125-6

Sengpiel, F., Blakemore, C., Kind, P., Harrad, R., 1994. Interocular suppression in the visual cortex of strabismic cats. J. Neurosci. 14, 6855–6871. https://doi.org/10.1523/JNEUROSCI.14-11-06855.1994

Sengpiel, F., Jirmann, K.-U., Vorobyov, V., Eysel, U.T., 2006. Strabismic suppression is mediated by inhibitory interactions in the primary visual cortex. Cerebral Cortex 16, 1750–1758.

Serences, J.T., 2004. A comparison of methods for characterizing the event-related BOLD timeseries in rapid fMRI. Neuroimage 21, 1690–1700.

Sereno, M.I., Dale, A.M., Reppas, J.B., Kwong, K.K., Belliveau, J.W., Brady, T.J., Rosen, B.R., Tootell, R.B.H., 1995. Borders of multiple visual areas in humans revealed by functional magnetic resonance imaging. Science 268, 889–893.

Sherman, S.M., Guillery, R.W., 2002. The role of the thalamus in the flow of information to the cortex. Philosophical Transactions of the Royal Society of London. Series B: Biological Sciences 357, 1695–1708.

Sincich, L.C., Park, K.F., Wohlgemuth, M.J., Horton, J.C., 2004. Bypassing V1: a direct geniculate input to area MT. Nat Neurosci 7, 1123–1128. https://doi.org/10.1038/nn1318

Takeuchi, H., Sekiguchi, A., Taki, Y., Yokoyama, S., Yomogida, Y., Komuro, N., Yamanouchi, T., Suzuki, S., Kawashima, R., 2010. Training of Working Memory Impacts Structural Connectivity. Journal of Neuroscience 30, 3297–3303. https://doi.org/10.1523/JNEUROSCI.4611-09.2010

Talairach, J., Tournoux, P., 1988. Co-planar Stereotaxic Atlas of the Human Brain: 3-dimensional Proportional System : an Approach to Cerebral Imaging. G. Thieme.

To, L., Thompson, B., Blum, J.R., Maehara, G., Hess, R.F., Cooperstock, J.R., 2011. A game platform for treatment of amblyopia. IEEE Transactions on Neural Systems and Rehabilitation Engineering 19, 280–289. https://doi.org/10.1109/TNSRE.2011.2115255

Tuna, A.R., Pinto, N., Brardo, F.M., Fernandes, A., Nunes, A.F., Pato, M.V., 2020. Transcranial magnetic stimulation in adults with amblyopia. Journal of Neuro-Ophthalmology 40, 185–192.

Vaina, L.M., Cowey, A., Jakab, M., Kikinis, R., 2005. Deficits of motion integration and segregation in patients with unilateral extrastriate lesions. Brain 128, 2134–2145.

Van Horn, S.C., Erişir, A., Sherman, S.M., 2000. Relative distribution of synapses in the A-laminae of the lateral geniculate nucleus of the cat. Journal of Comparative Neurology 416, 509–520. https://doi.org/10.1002/(SICI)1096-9861(20000124)416:4<509::AID-CNE7>3.0.CO;2-H

Warner, C.E., Goldshmit, Y., Bourne, J.A., 2010. Retinal afferents synapse with relay cells targeting the middle temporal area in the pulvinar and lateral geniculate nuclei. Frontiers in neuroanatomy 4, 8.

Xu, J.P., He, Z.J., Ooi, T.L., 2012. Perceptual learning to reduce sensory eye dominance beyond the focus of top-down visual attention. Vision Research 61, 39–47.

Xu, J.P., He, Z.J., Ooi, T.L., 2010. Effectively reducing sensory eye dominance with a push-pull perceptual learning protocol. Current Biology 20, 1864–1868.

Xue, J.T., Ramoa, A.S., Carney, T., Freeman, R.D., 1987. Binocular interaction in the dorsal lateral geniculate nucleus of the cat. Experimental brain research 68, 305– 310. https://doi.org/10.1007/BF00248796

Yang, E., Blake, R., McDonald, J.E., 2010. A new interocular suppression technique for measuring sensory eye dominance. Investigative Ophthalmology & Visual Science 51, 588–593. https://doi.org/10.1167/iovs.08-3076

Zhang, J.-Y., Cong, L.-J., Klein, S.A., Levi, D.M., Yu, C., 2014. Perceptual learning improves adult amblyopic vision through rule-based cognitive compensation. Investigative ophthalmology & visual science 55, 2020–2030.

Zhang, P., Bobier, W., Thompson, B., Hess, R.F., 2011. Binocular balance in normal vision and its modulation by mean luminance. Optometry and Vision Science 88, 1072–1079.

Zhou, J., Huang, P.-C., Hess, R.F., 2013. Interocular suppression in amblyopia for global orientation processing. Journal of vision 13, 19–19. https://doi.org/10.1167/13.5.19

